# Transcript degradation and codon usage regulate gene expression in a lytic phage

**DOI:** 10.1101/647024

**Authors:** Benjamin R. Jack, Daniel R. Boutz, Matthew L. Paff, Bartram L. Smith, Claus O. Wilke

## Abstract

Many viral genomes are small, containing only single- or double-digit numbers of genes and relatively few regulatory elements. Yet viruses successfully execute complex regulatory programs as they take over their host cells. Here, we propose that some viruses regulate gene expression via a carefully balanced interplay between transcription, translation, and transcript degradation. As our model system, we employ bacteriophage T7, whose genome of approximately 60 genes is well annotated and for which there is a long history of computational models of gene regulation. We expand upon prior modeling work by implementing a stochastic gene expression simulator that tracks individual transcripts, polymerases, ribosomes, and RNases participating in the transcription, translation, and transcript-degradation processes occurring during a T7 infection. By combining this detailed mechanistic modeling of a phage infection with high throughput gene expression measurements of several strains of bacteriophage T7, evolved and engineered, we can show that both the dynamic interplay between transcription and transcript degradation, and between these two processes and translation, appear to be critical components of T7 gene regulation. Our results point to a generic gene regulation strategy that may have evolved in many other viruses. Further, our results suggest that detailed mechanistic modeling may uncover the biological mechanisms at work in both evolved and engineered virus variants.

## Introduction

Bacteriophages are widely established model systems in comparative genomics [1], experimental evolution [2], and synthetic biology [3]. Their rapid replication rates, ease of culturing, and moderately small genome sizes make them ideal candidates for studies into the evolution of both natural and engineered variants, including the adaptation of variants with rearranged genomes [4] or modified codon usage [5], parallel evolution [6, 7], adaptation to nonstandard genetic codes [8], or the structural effects of adaptive mutations [9]. One of the most widely studied bacteriophages is bacteriophage T7, whose genome was first sequenced in 1983 [10]. A wealth of specific knowledge about T7’s genes, gene regulation, and gene interactions has been accumulated since [11], and this knowledge has enabled the development of detailed mechanistic models of the viral life cycle inside a bacterial cell [12–16].

Bacteriophage T7 infects *E. coli* and rapidly lyses the cell, in as little as 11 minutes at 37°C [10,17]. In this 11 minute time period, the phage produces 20-40 viable virions from a single *E. coli* cell [18]. To produce these virions, the phage must coordinate and assemble nearly 10,000 copies of the major capsid protein alone (415 per virion) [10]. In accomplishing this high output, T7 generates stable transcripts and has a genome whose codon usage is optimized for its *E. coli* host [5]. Given how quickly T7 replicates, one might assume that transcription and translation would be tightly coupled to produce the correct ratios of phage proteins. In contrast to this expectation, however, T7 RNA polymerase (the source of most phage transcripts), moves at an order of magnitude faster than the translation machinery [19–21]. Thus, the balance between transcription and translation is not obvious. We ask here how T7 regulates its genes to ensure appropriate relative ratios of transcripts and proteins. We address this question by combining a detailed mechanistic model of the viral life cycle with high throughput gene expression data obtained from evolved and engineered T7 strains.

We specifically test the hypothesis that a key component of gene regulation in phage T7 is targeted degradation. We analyze RNA-sequencing data from T7 infections [18,22], combined with proteomics data where available, and we test various gene regulation mechanisms via a detailed computational simulation of a phage infection. We detect and analyze a region of the T7 genome that is down-regulated despite an absence of transcriptional terminators. We then propose and computationally test a directional degradation mechanism that may explain this pattern of down-regulation. Next, we explore the relationship between transcripts and proteins during the course of a simulated T7 infection. Lastly, we assess the interaction between codon usage and promoter strength in the production of the most abundant phage protein, the major capsid protein. In aggregate, we argue that T7 gene regulation is controlled primarily via the dynamic interplay of transcription, transcript degradation, and translation.

## Results

### A brief introduction to T7 biology

The bacteriophage T7 genome contains nearly 60 genes, divided into three classes [10,11]. Class I consists of genes transcribed by the host RNA polymerase, including T7 RNA polymerase (RNAP). Classes II and III are transcribed by T7 RNAP: class II encodes DNA polymerase and proteins associated with genome replication, and class III encodes structural proteins.

Genes are numbered in the order in which they are encoded in the genome, and all genes are encoded on the same strand with minimal overlaps [10]. The phage genome contains 17 promoters recognized by T7 RNAP. A single T7 terminator T*ϕ* is located immediately after the major capsid gene, gene *10*. In addition to these regulatory elements affecting transcription, there are 10 known RNase cleavage sites. Because of its genomic architecture, T7 produces many polycistronic transcripts of varying lengths.

T7 infections proceed rapidly. At 30 °C, T7 wild type lyses *E. coli* cells in approximately 30 minutes [10]. At higher temperatures and with laboratory adapted strains, lysis occurs even more rapidly, as quickly as 11 minutes post infection at 37 °C [17]. During the course of infection, T7 shuts down all *E. coli* gene expression and degrades both the *E. coli* genome and its transcripts [11,18].

In this work, we consider five different strains of T7. The bulk of our analysis is performed on strain T7_61_, a wild-type strain adapted for 20 hours to grow under laboratory conditions at 37 °C [5,23]. This strain lyses *E. coli* within 11 minutes. Unless otherwise specified, “T7” or “T7 wild type” refers to this strain throughout this work. To make a comparison to a diverged strain, we additionally collected RNA abundances for the progenitor of T7_61_, called T7^+^ [24], grown at 30 °C. T7_61_ differs from T7^+^ in a deletion of nearly 1500 bases near the beginning of the genome and in several point mutations [5,23]. We also considered three engineered variants of T7_61_, one with gene *10* codon-deoptimized [5], one with the two promoters *ϕ*9 and *ϕ*10 upstream of gene *10* knocked out [22], and one in which both the codon-deoptimization and the promoter knockouts have been applied [22].

### Transcripts degrade during infection

Because of T7’s short infection cycle and the stability of its mRNA transcripts *in vitro,* prior studies assumed that the effect of degradation on T7 gene expression was negligible or at least uniform [10,12,25]. If no transcripts degraded during infection, we would expect the distribution of transcript abundances late in the infection cycle to have two specific features. First, for the genes transcribed ahead of the single terminator located at the end of gene *10,* downstream genes should have higher transcript abundances than upstream genes, due to the multiple promoters. Second, transcript abundances of genes downstream of the terminator (i.e., downstream of gene *10*) should be lower than those of upstream genes. To assess whether T7 transcript abundances match these expectations, we reanalyzed RNA-sequencing data from a previous study that had collected T7 RNA abundances at 1, 5, and 9 minutes post infection [18].

At 1 min post infection, we detected few T7 genes [18]. However, all T7 genes were expressed at 5 and at 9 min (Fig. 1). We found that transcript abundances at 5 min were approximately uniform but noisy, except for a clear drop downstream of the terminator T*ϕ* (Fig. 1A). By contrast, at 9 min, expression had shifted from class II to class III genes (Fig. 1B) and displayed much more systematic variation. In particular, we observed a cluster of class II genes between genes *3.8* and *6.5* that had expression levels lower than that of upstream genes. This region contains no known terminators. If a terminator were present, we would expect to see a similarly down-regulated region of genes at 5 min post infection. However, we did not detect this under-expressed cluster at 5 min (Fig. 1B). Moreover, throughout the class II genes, we observed relative increases in expression of downstream genes where no promoters were present. In aggregate, these observations are inconsistent with a model of T7 gene regulation in which transcripts do not degrade, or in which degradation is uniform.

**Figure 1:**
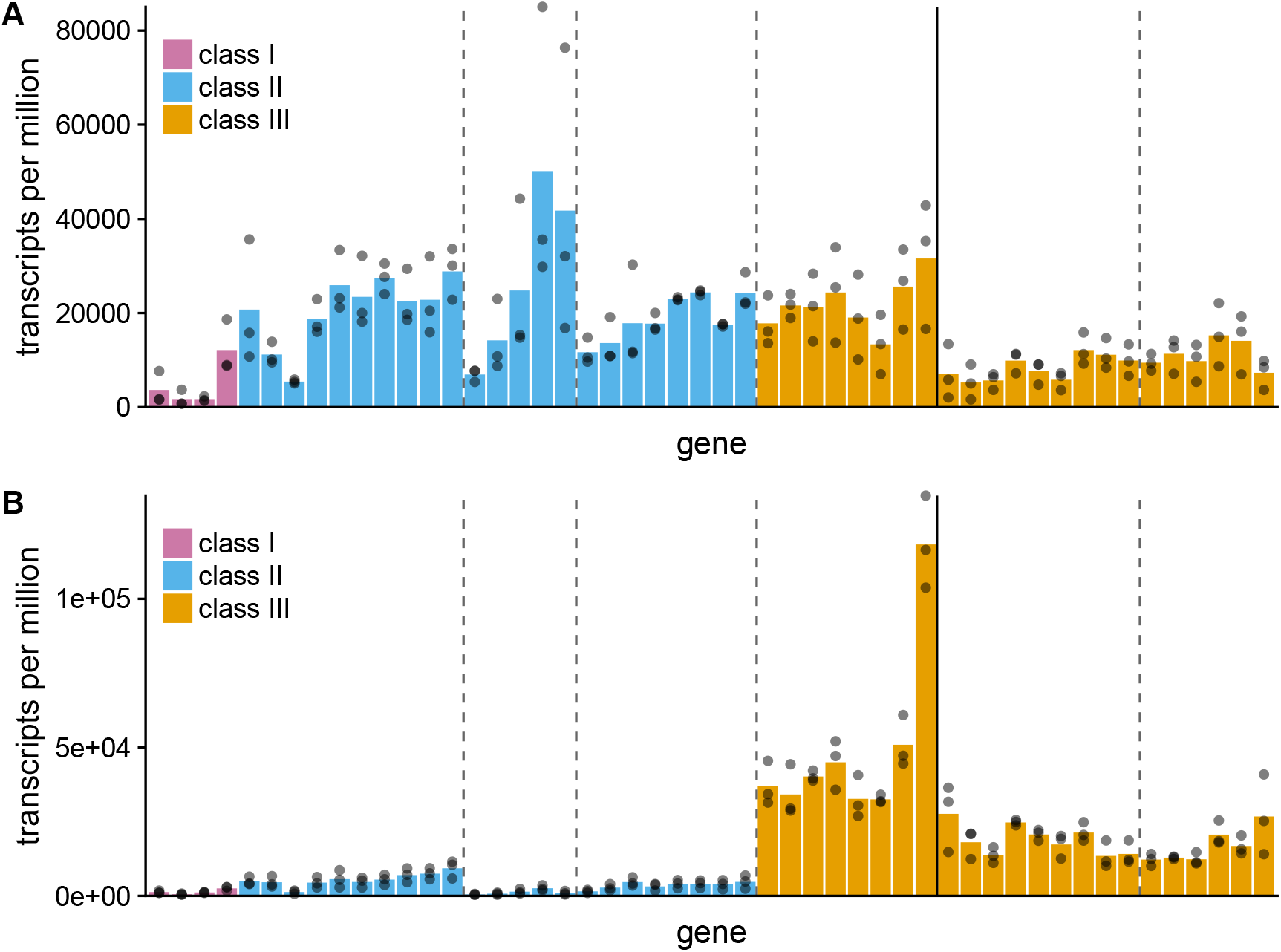
Relative transcript abundances across the T7 genome at 5 minutes and 9 minutes post infection. Each colored bar represents one gene, and genes are arranged from left to right in the order in which they appear in the T7 genome. Dashed vertical lines indicate RNAse cleavage sites R3.8, R4.7, R6.5, and R13, respectively. Solid vertical lines indicate the terminator T*ϕ*. (A) Gene expression at 5 minutes post infection. Classes II and III show similar patterns of expression. (B) Gene expression at 9 minutes post infection, just before lysis. Class III show higher expression compared to class II. Transcript abundance levels decrease between genes *3.8* and *6.5.* No terminators are present in this genomic region, suggesting that extensive transcript degradation produces the sudden drop-off in transcript abundance.

To further characterize this cluster of unexpectedly down-regulated class II genes, we examined individually mapped reads in the T7 genome. We found that across the whole genome, raw read counts followed the same broad pattern as transcript abundances (Fig. 2A). The highest read counts fell within gene *10*, and counts decreased after the terminator T*ϕ*. Again, we observed the down-regulation of a cluster of class II genes. This down-regulated region contains several regulatory elements, including both RNase cleavage sites and promoters. We examined four regulatory elements in this region, R3.8/*ϕ*3.8 (Fig. 2B) and R6.5/*ϕ*6.5 (Fig. 2C). In each case, the transcription start site lies upstream of the RNase cleavage site. In the R3.8 region we saw a sustained decrease in read counts (at least 500bp downstream), but in the R6.5 region we found the opposite trend. Read counts recovered in fewer than 500bp. We concluded that transcript synthesis alone could not explain these observations, and that RNase cleavage sites may contribute to transcript degradation.

**Figure 2:**
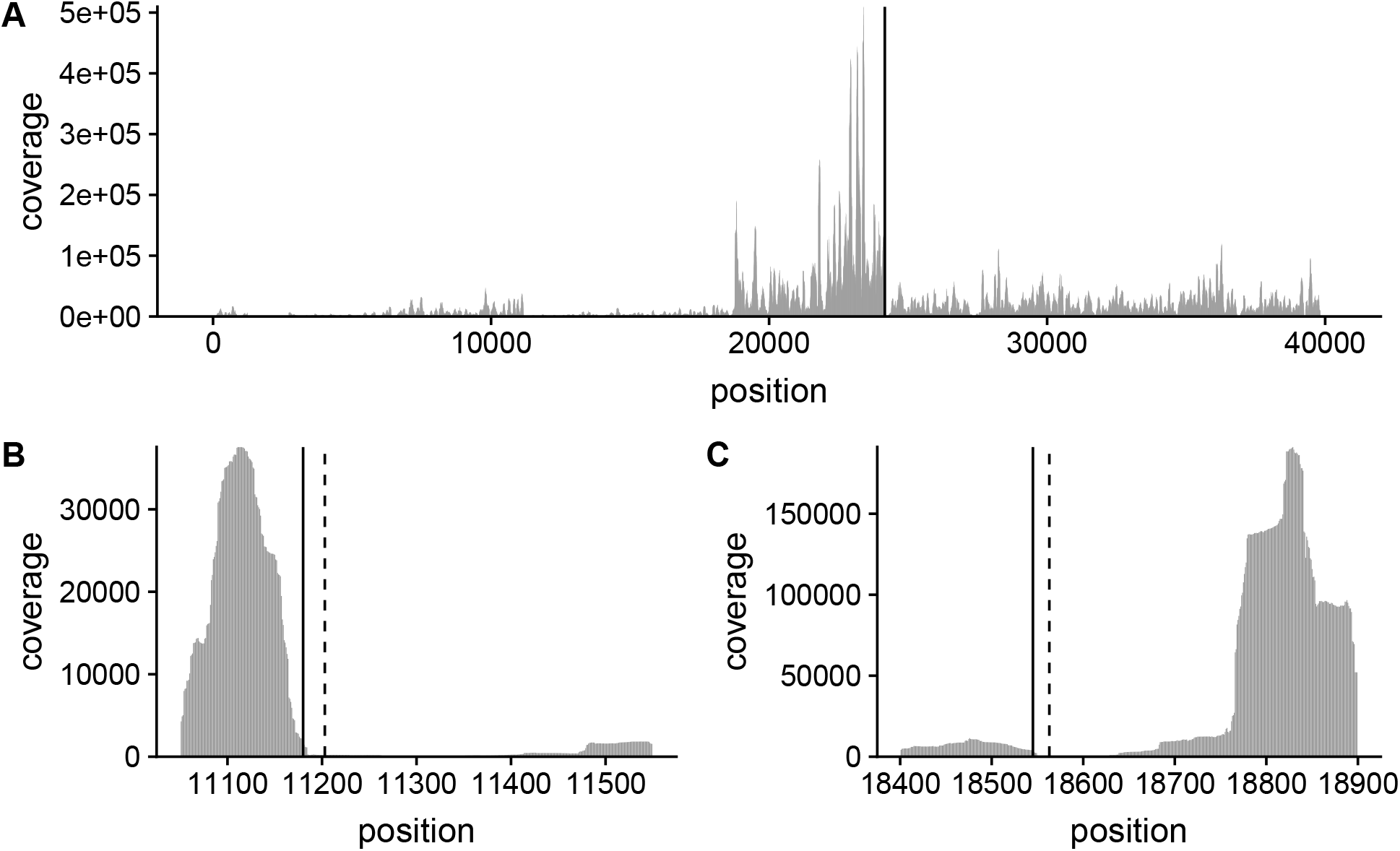
RNA-sequencing genome coverage map of bacteriophage T7, 9 minutes after infection. Each bar represents the number of reads that map to that specific position in the genome. (A) A coverage map of the entire 40kb T7 genome. The solid vertical line represents T*ϕ*, the terminator located downstream of gene *10.* Mapped reads decrease downstream of this terminator. (B) The region of the genome surrounding RNase cleavage site R3.8. Mapped reads decrease downstream of the cleavage site. The solid vertical line represents the transcription start site from promoter *ϕ*3.8 and the dashed vertical line represents the cleavage site. (C) The region of the genome surrounding RNase cleavage site R6.5. Here, mapped reads sharply increase downstream of *ϕ*6.5 and R6.5. The solid vertical line represents the transcription start site for *ϕ*6.5 and the dashed vertical line represents the cleavage site.

Lastly, we verified the generality of down-regulation in class II genes by conducting experiments at 30 °C, using the progenitor wild type strain T7^+^. Our aim was to determine if the degradation patterns we observed in the lab-adapted strain were unique to that strain or instead represented a general feature of T7 biology. Since lysis occurs after 25-30 minutes at 30 °C, we collected samples at 5, 10, 15, 20, and 25 minutes. We then compared an early time point (10 min) to a later time point (25 min). We found the same expression patterns within class II as we had observed for the lab-adapted strain (Fig. 3). Outside of class II expression, we observed one major difference in gene *19.5* expression, which was elevated in T7^+^ but not in the lab-adapted T7_61_. This gene has unknown function and we do not know why it is so highly expressed in T7^+^. Overall, however, we found that the evidence for differential transcript degradation is consistent across multiple strains of T7 grown under different conditions. Consequently, this degradation pattern is likely a general feature of T7 gene regulation.

**Figure 3:**
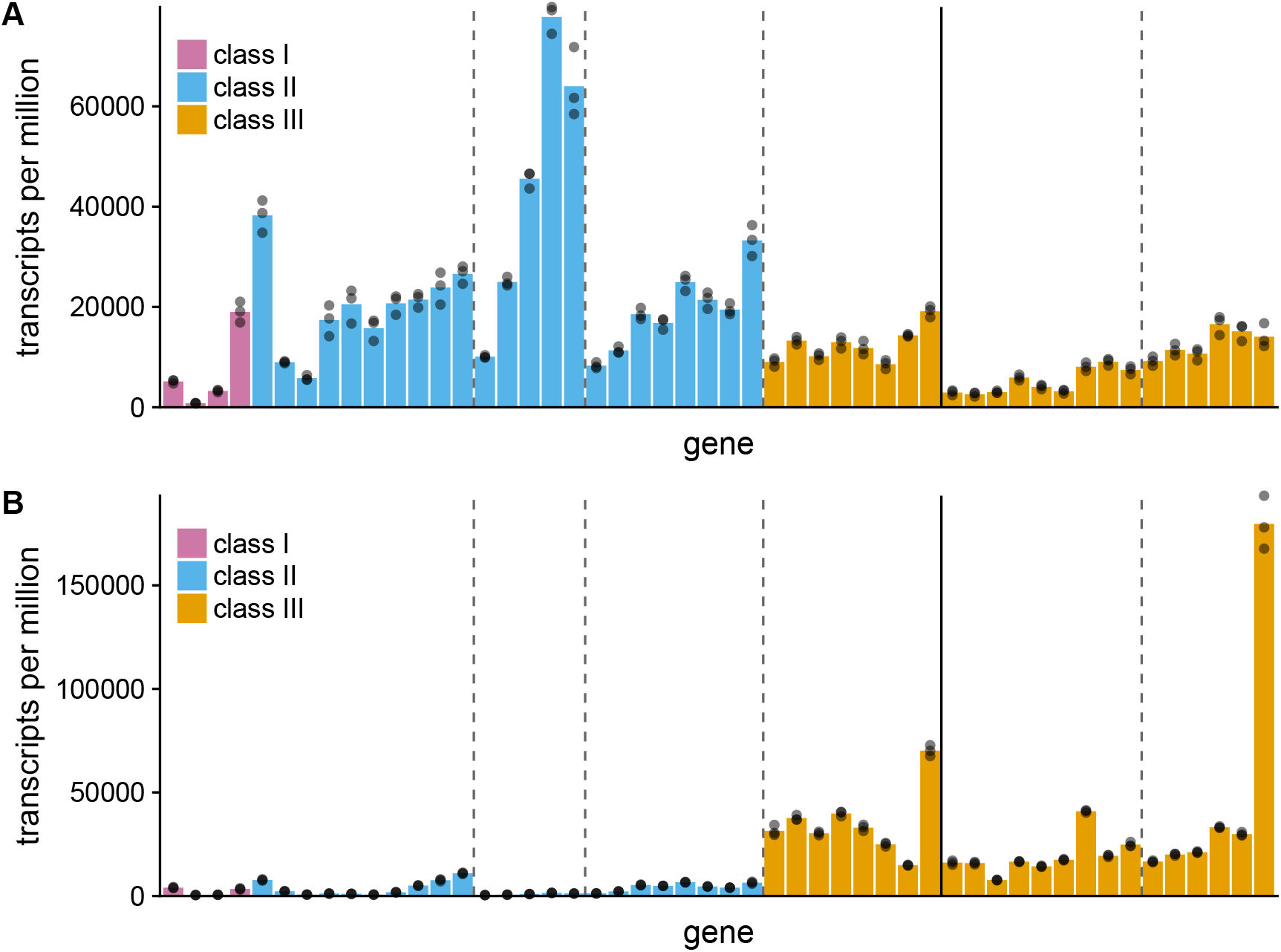
Relative transcript abundances across the T7 genome from T7^+^, a strain that has not been adapted to laboratory conditions. Samples were taken at 10 minutes and 25 minutes post infection and at 30 °C (compared to 37 °C in prior experiments). Each colored bar represents one gene, and genes are arranged from left to right in the order in which they appear in the T7 genome. Dashed vertical lines indicate RNAse cleavage sites R3.8, R4.7, R6.5, and R13, respectively. Solid vertical lines indicate the terminator T*ϕ*. (A) Gene expression at 10 minutes post infection. Classes I and II show higher expression levels than class III. (B) Gene expression at 25 minutes post infection, just before lysis. Class III shows higher expression compared to class II. Gene *19.5* has the highest overall expression. Transcript abundance levels decrease between genes *3.8* and *6.5.* No terminators are present after gene *3.8,* suggesting that extensive transcript degradation produces the sudden drop-off in transcript abundance. Although cultured at a different temperature and with a different strain of T7, both the 30 °C and the 37 °C experiment show the same region of down-regulated class II genes.

### Degradation can produce gene expression patterns similar to that of promoters and terminators

Motivated by the preliminary evidence for transcript degradation in T7, we next employed a simulation model to assess how promoters, terminators, and transcript degradation processes can interact to determine gene expression over time. The simulation software we used, Pinetree, simulates prokaryotic gene expression with single-nucleotide resolution [26]. Pinetree tracks individual polymerases on DNA and ribosomes on RNA, and it supports polycistronic transcripts and a variety of competing regulatory mechanisms including promoter binding, termination, transcript degradation, and variable translation rates due to codon usage.

Pinetree employs a directional model of transcript degradation modeled after observations from *E. coli* [27–29]. In *E. coli,* degradation occurs either from the 5’-end of a newly-synthesized transcript or from the 5’-end of a transcript recently cleaved by an RNase III ribnuclease [30]. A combination of endo- and exonucleases degrade transcripts, but the net effect is that transcript degradation is directional—the 5’ end of a transcript has a shorter lifespan than the 3’ end [31]. In Pinetree, this degradation is simulated via RNAses that bind to the 5’-end of transcripts and degrade in the 5’-to-3’ direction. While no such 5’-to-3’ RNase actually exists in *E. coli,* it approximates the joint effect of several endo- and exo-nucleases that collectively tend to degrade the 5’-end of the transcript more quickly than the 3’-end [31]. Moreover, ribosomes compete with RNAses for access to transcripts [31]. Pinetree captures this competition between ribosomes and RNAses, and it also implements the difference in degradation rate between newly-synthesized and recently cleaved transcripts [32].

To test how the interplay between transcription and degradation affects gene regulation, we first studied a simple toy model. Our aim was to account for two observations in T7: i) increases in downstream transcript abundances in the absence of promoters, and ii) reduced transcript abundances in downstream genes in the absence of terminators. We constructed, *in silico,* a linear plasmid containing three genes of equal length, for three proteins X, Y, and Z. The plasmid contained a single promoter upstream of gene *X* and a single terminator after gene *Z* (Fig. 4A). We simulated gene expression for 240 seconds in an *E. coli*-like cellular environment, with a fixed pool of RNA polymerases. We observed at all time points that transcript abundances were greatest for gene Z, followed by gene *Y* and then gene *X* (Fig. 4B, C, D). Since the plasmid contained one promoter and one terminator, the simulation produced only tricistronic transcripts. However, since transcripts degraded directionally, gene *X* had the lowest expression level, and expression levels increased from gene *X* to *Y* to *Z* (Fig. 4B, C). These simulation results show that polycistronic transcripts with directional degradation are sufficient to produce gene expression patterns that mimic the effects of promoters.

**Figure 4:**
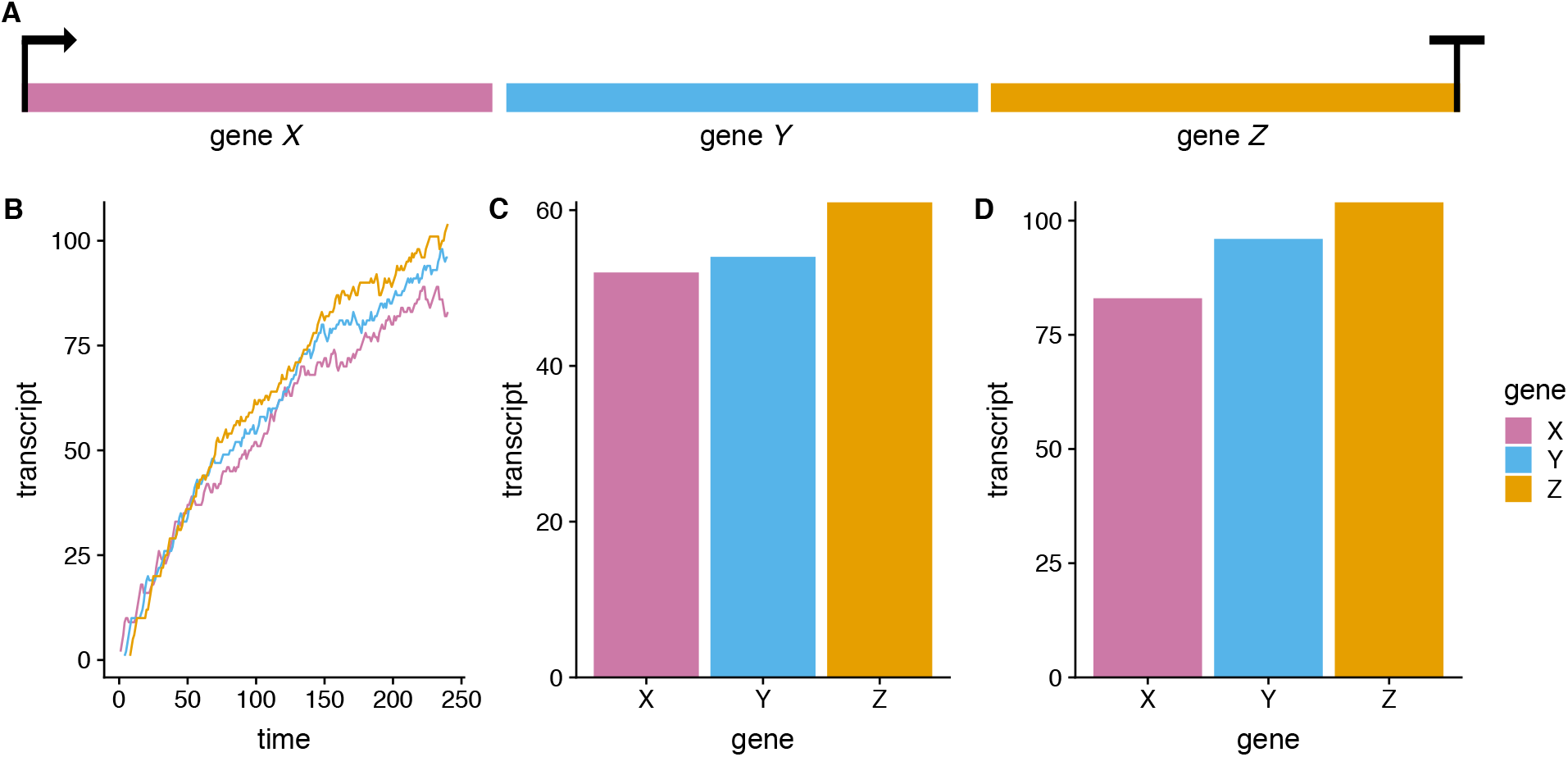
Simulation of transcription and transcript degradation for a plasmid containing three genes of equal length. (A) The plasmid contains a single promoter upstream of gene X, generating polycistronic transcripts that contain all three genes. RNAses degrade transcripts from the 5’-to-3’ direction. (B) Transcript abundances over time during a 240s simulation. (C) Transcript abundances at 100s. (D) Transcript abundances at 240s. As the simulation progresses, directional degradation reduces abundances of genes encoded closer to the 5’-end of the transcript.

We next introduced an RNase cleavage site and an additional promoter upstream of gene *Z* into the linear plasmid simulation (Fig. 5A). This arrangement of RNase cleavage site and promoter is common in the T7 genome [10]. Adding these two regulatory elements created a dynamic gene expression pattern in which earlier time points showed the expected ramp of increased gene expression from genes *X* and *Y* to *Z* (Fig. 5B, C). Later time points, however, showed a different pattern: gene *Z* transcripts had lower abundance than did transcripts of genes *X* and *Y* (Fig. 5B, D). Thus, the addition of RNase cleavage sites to the simulation was sufficient for recreating gene expression patterns that mimic terminators at later time points but not at earlier time points. This trend in gene expression is similar to the trend observed in experimental transcript data of T7.

**Figure 5:**
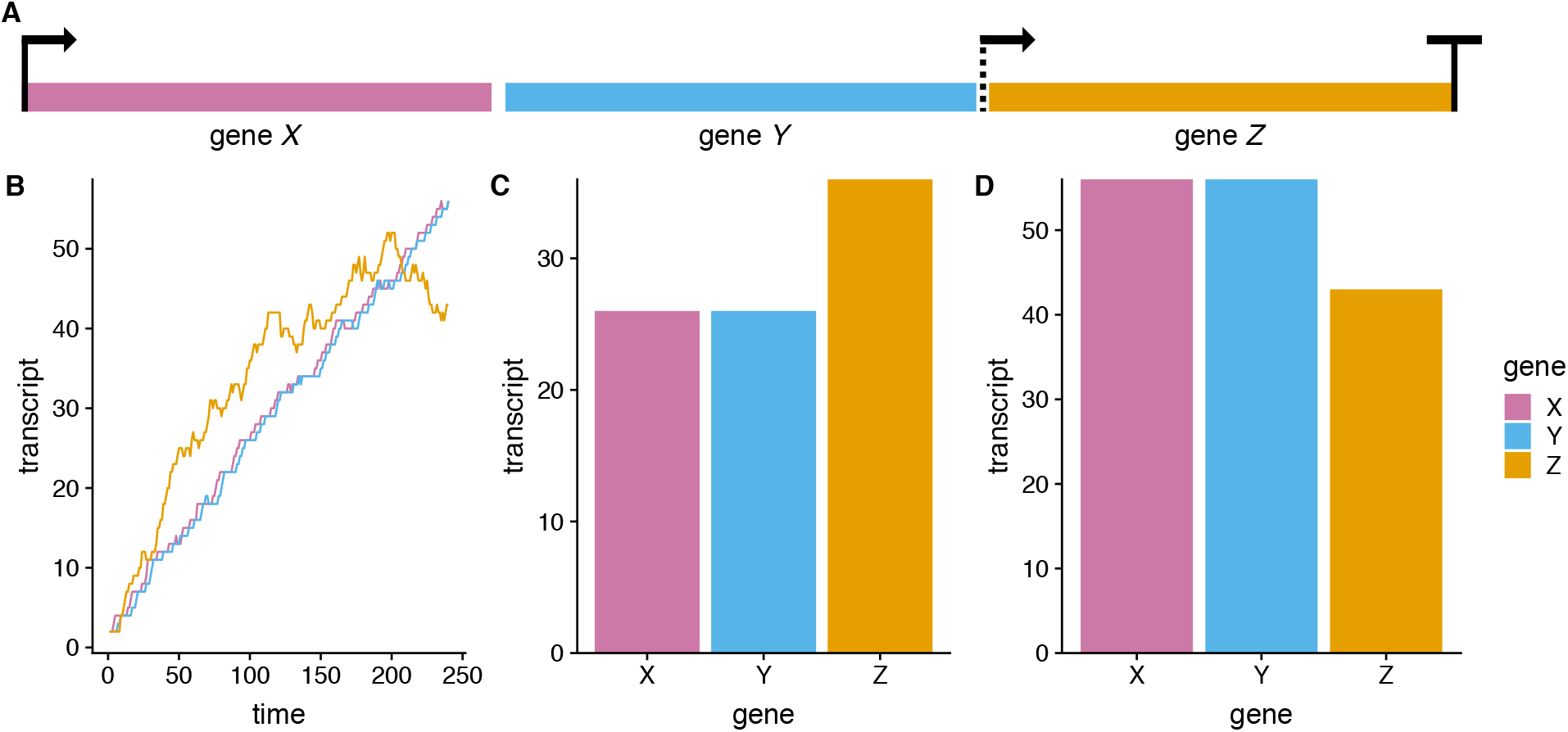
Simulation of gene expression for a three-gene plasmid containing two promoters and an RNase cleavage site. (A) Promoters are encoded upstream of genes *X* and *Z* (represented by arrows). An RNase cleavage site (dashed line) is encoded upstream of gene *Z*, and downstream of the promoter (arrow). Degradation proceeds in the 5’-to-3’ direction from the 5’-end of the transcript and from the RNase cleavage site. (B) Transcript abundances over time during a 240s simulation. Transcript abundances for gene *Z* are higher than other genes initially, but over time gene *Z* transcripts degrade more quickly than gene *Y* and gene *Y* transcripts become most abundant. (C) Transcript abundances at 100s. (D) Transcript abundances at 240s. An internal RNase cleavage site near a promoter in the plasmid creates dynamic gene expression patterns. Initially, the stronger promoter upstream of *Z* creates more transcripts than *X* and *Y*, until it begins to reach equilibrium with degradation due to the RNase cleavage site. Degradation at the cleavage site is stronger than at the 5’ end of the transcript, so transcripts of *X* and *Y* continue to increase to abundances higher than that of *Z*.

### Degradation produces transcript ramps and cliffs in simulations of phage T7

After demonstrating that degradation and RNase cleavage sites are sufficient for creating dynamic gene expression patterns in a three-gene linear plasmid, we next considered whether a simulation of the full T7 genome would reveal similar expression patterns. Again, we used the Pinetree simulator, now to simulate the full T7 genome both with and without RNase cleavage sites and degradation. We attempted to represent the T7 genome as accurately as possible, matching the genetic architecture of the reference sequence [10]. We simulated T7 among cellular resources representative of an *E. coli* cell. These resources included RNA polymerases, ribosomes, and RNases, as well as secondary reactions between synthesized T7 proteins (Methods). However, we did not explicitly simulate expression of any *E. coli* genes. The simulation reliably reproduced the overall patterns of gene expression seen in T7: Class I genes are expressed the earliest but reach on average the lowest transcript abundances, and class III genes are expressed the latest and reach on average the highest transcript abundances (Fig. 6).

**Figure 6:**
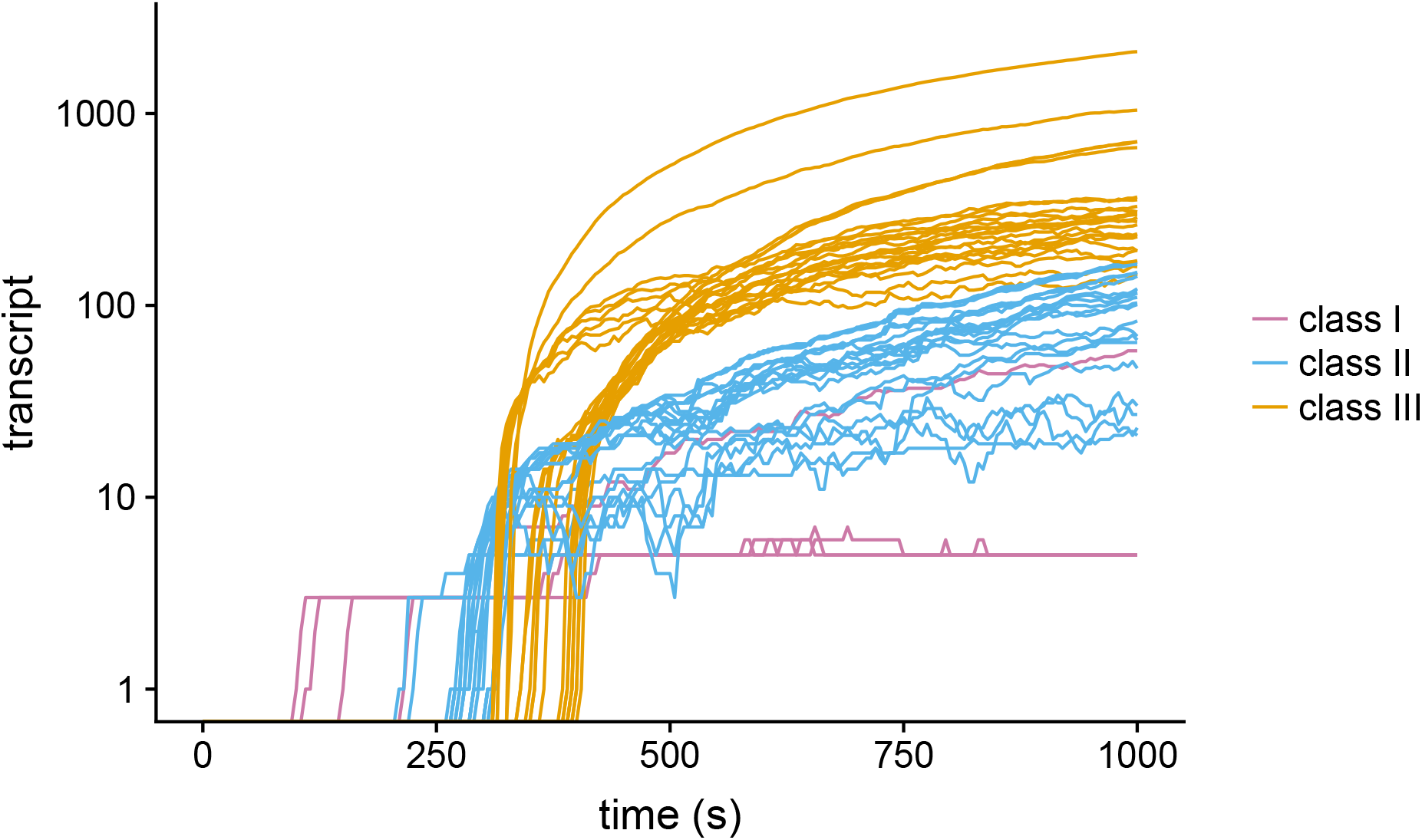
Transcript abundances show differential gene expression over time in a simulated T7 infection. The simulation includes transcript degradation and RNase cleavage sites. Class I genes are expressed first, followed by class II and then class III genes.

We next analyzed how measured transcript abundances for individual transcripts late in the infection (Fig. 7A) compared to their simulated counterparts (Fig. 7B, C). When simulating T7 with neither RNase cleavage sites nor degradation, we found that the simulation captured the broad expected patterns of gene expression for the three classes of T7 genes (Fig. 7B). Transcript abundances of downstream genes were higher than those of upstream genes, except after the terminator T*ϕ*. However, expression in the region of class II genes between *3.8* and *6.5* differed from expression in our experimental data (Fig. 7A). In our experimental data, we also observed increases in downstream gene expression where no promoters are present, which our simulation without degradation did not capture. We refer to these gradually increasing downstream expression patterns as *ramps,* and the decrease in downstream expression as *cliffs.*

**Figure 7:**
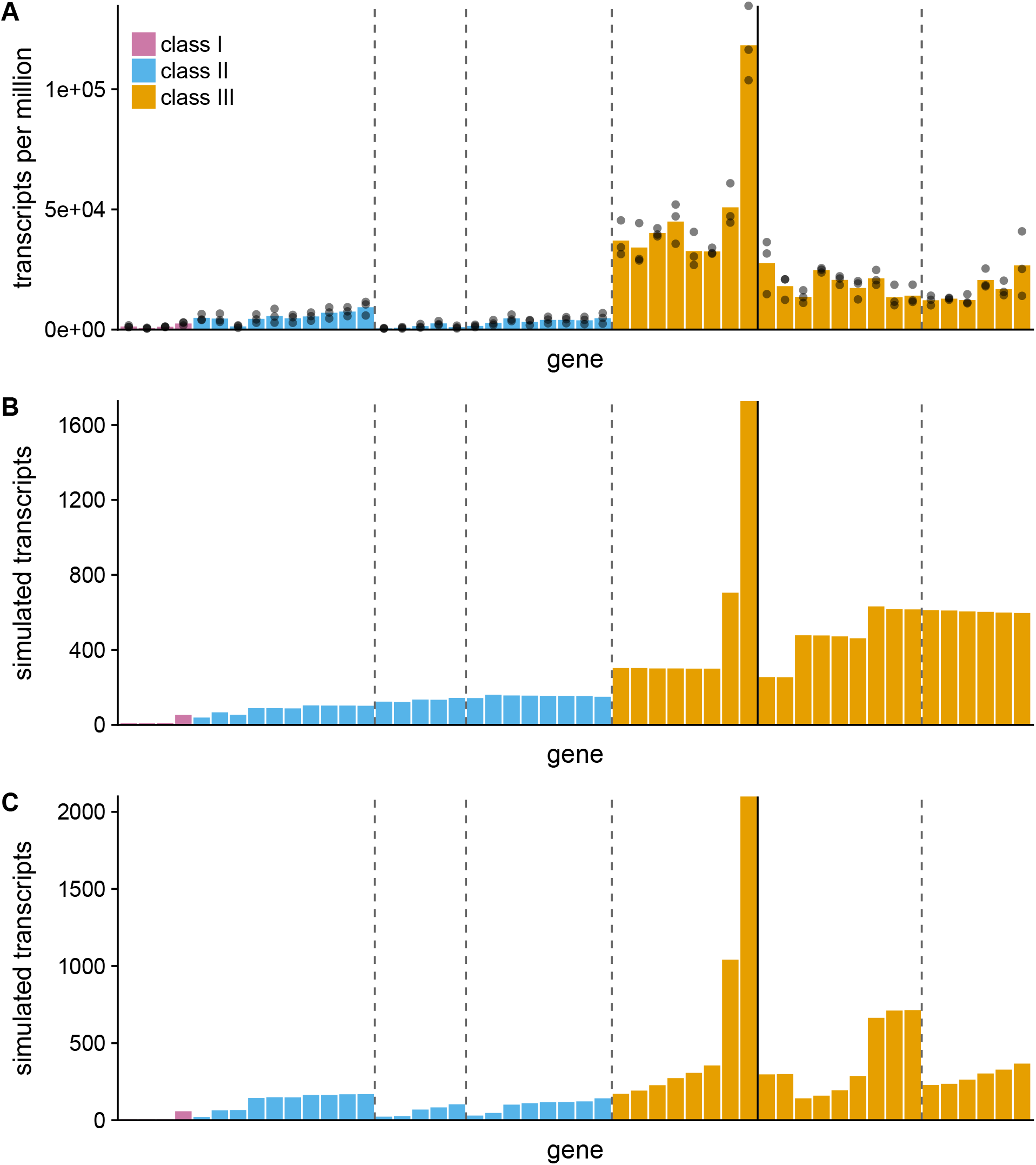
Simulations of T7 that include internal RNase cleavage sites and transcript degradation create transcript abundance distributions that resemble experimental distributions. Each colored bar represents one gene, and genes are arranged from left to right in the order in which they appear in the T7 genome. The solid vertical line represents the terminator T*ϕ* and the dashed lines represent RNase cleavage sites R3.8, R4.7, R6.5, and R13, respectively. (A) Distribution of experimental transcript abundances of bacteriophage T7, 9 minutes after infection. (B) Distribution of simulated transcript abundances, 1000s simulation time after infection, without RNase cleavage sites or degradation. We observe no reduction in gene expression between genes *3.8* and *6.5.* (C) Distribution of simulated transcript abundances, 1000s simulation time after infection, from a simulation that includes RNAse cleavage sites and transcript degradation. The region between R3.8 and R6.5 shows lower transcript abundances than are seen for upstream genes. This expression pattern is similar to that of the experimental observations. Including directional degradation and RNase cleavage sites in a simulation of T7 are sufficient to reproduce patterns of reduced gene expression between genes *3.8* and *6.5* from experimental data.

To test whether transcript degradation and RNase cleavage sites were sufficient to explain expression ramps and cliffs, we conducted a set of simulations that included the 10 RNase cleavage sites in the T7 genome and directional transcript degradation (Fig. 7C). In this simulation, we observed gene expression ramps and cliffs after RNase cleavage sites. Simulated transcript degradation and RNase cleavage sites created a distribution of transcript abundances more qualitatively similar to our experimental distributions than a model without degradation.

### Relationship between transcript and protein abundances differs among gene classes

We also considered the relationship between transcript and protein abundances. T7 RNA polymerase moves at approximately 230bp/s, while ribosomes only translate at a rate of 30bp/s [33]. This speed difference means that translation significantly lags transcription, and that transcription and translation are likely uncoupled in T7 [34]. We assessed this hypothesis by examining the relationship between both protein and transcript abundances during the course of infection, in experiments and in simulations. Our aim was to determine how changes in RNA abundances affect protein abundance.

We compared RNA and protein abundances at 5 minutes and at 9 minutes post infection in the lab-adapted T7 wildtype grown at 37 °C (Fig. 8A). We found that at 5 min, class III genes showed a stronger correlation between RNA and protein abundances than did class II genes (Pearson’s *r*; class II genes: *r* = 0.134, *p* = 0.584; class III genes: *r* = 0.628, *p* = 0.00530), and that class II and class III genes clustered together in transcript-protein space. At 9 minutes, class II and class III expression separated along the transcript axis, but not along the protein axis (Fig. 8A). Class III genes continued to show a stronger correlation between transcript and protein abundances than did class II genes (Pearson’s r; class II genes: *r* = 0.245, *p* = 0.299; class III genes: *r* = 0.700, *p* = 0.000596). This result suggested that either class II transcripts degraded or class III transcripts increased between 5 and 9 minutes, and that this change in transcript abundances was uncoupled from protein production.

**Figure 8:**
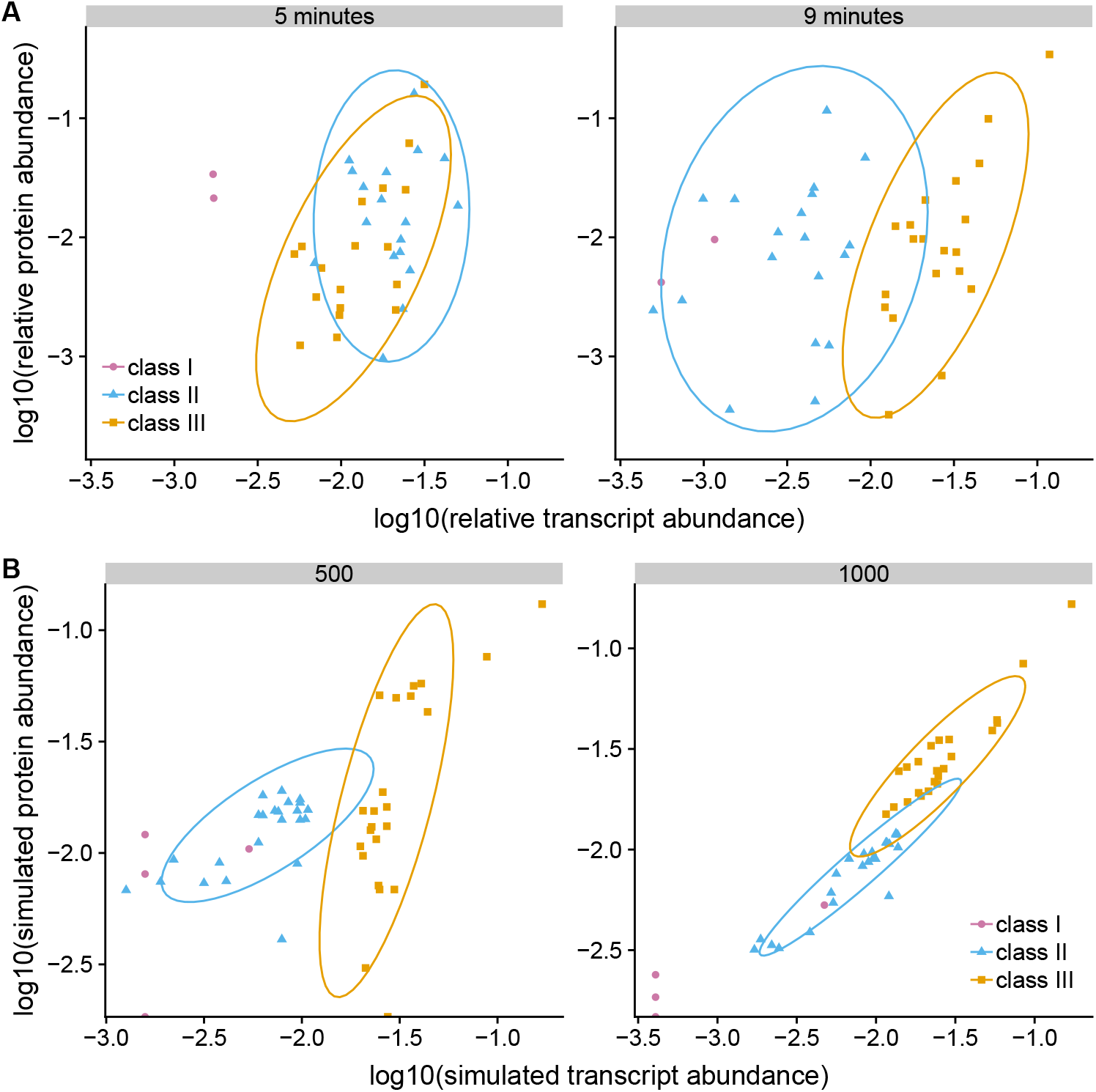
The relationship between protein and transcript abundances changes during the course of a T7 infection, both in experiments and in simulations. (A) Protein and transcript abundances from experiments at 5 and 9 minutes after infection. Correlations between protein and transcripts change between 5 minutes (Pearson’s *r*; class II genes: *r* = 0.134, *p* = 0.584; class III genes: *r* = 0.628, *p* = 0.00530) and 9 minutes (class II genes: *r* = 0.245, *p* = 0.299; class III genes: *r* = 0.700, *p* = 0.000596). (B) Simulated protein and transcript abundances at 500s (Pearson’s *r*; class II genes: *r* = 0.596, *p* = 2.10 × 10^-3^; class III genes: *r* = 0.750, *p* = 5.80 × 10^-5^) and 1000s (class II genes: *r* = 0.937, *p* = 1.58 × 10^-11^; class III genes: *r* = 0.926, *p* = 2.26 × 10^-10^) after infection. As in the experimental distributions, we observe two distinct classes of gene expression. The simulation captures changes in the relationship between protein, but shows proteins and transcripts as more strongly correlated than they are in the experimental observations. In the simulations, either transcript abundances are too high or protein abundances too low relative to the experimental observations.

Simulations of T7 gene expression yielded correlations of transcripts and proteins within classes II and III both early (Pearson’s *r*; class II genes: *r* = 0.596, *p* = 2.10 × 10^-3^; class III genes: *r* = 0.750, *p* = 5.80 × 10^-5^) and late in the infection (Pearson’s *r*; class II genes: *r* = 0.937, *p* = 1.58 × 10^-11^; class III genes: *r* = 0.926, *p* = 2.26 × 10^-10^) (Fig. 8B). Overall, correlations were too strong in the simulations compared to the experimental data. This finding suggested that either transcript degradation was too strong in our simulations, creating a tight coupling of RNA and protein abundances, in particular late in the infection, or that we were not properly accounting for dynamics in translation.

One important component of translation we did not model is dynamic tRNA pools. In our simulation, we assume that the availability of charged tRNAs never explicitly limits translation. Some codons are always translated more quickly than others. However, in living *E. coli* cells, abundances of charged tRNAs may change during infection, due to the stress the cells experience during infection, and such changes would affect codon-specific translation rates [35,36]. Thus, further work may be needed to extend Pinetree to include a realistic translation model.

### Promoter knockouts and codon deoptimization have antagonistic effects on gene expression

Finally, we wanted to assess to what extent our T7 simulation generalizes to more complex genome modifications. We considered two specific modifications for which we had existing experimental data, codon deoptimization and promoter knockouts [18,22]. Both of these modifications reduce viral fitness and have been proposed as viable approaches to viral attenuation. Codon deoptimization is the process of replacing common codons (relative to the *E. coli* host genome) with rare codons, to reduce translation rates. It has been applied to T7 gene *10* [5,18]. Gene *10* encodes the major capsid protein, the most abundantly expressed phage protein, and reducing its abundance is expected to reduce phage fitness. Similarly, because of T7’s genome architecture containing many overlapping open reading frames, it is possible to attenuate but not kill the virus by knocking out key promoters. Prior work has considered the effects of knocking out the promoters upstream of genes *9* and *10* (*ϕ*9 and *ϕ*10), individually and in combination, and also in combination with codon deoptimization of gene *10* [22].

Reducing the expression of gene *10* by either codon deoptimization or promoter knockout resulted in significant fitness reduction [5,22]. Codon deoptimization resulted primarily in a reduction in protein 10 abundance [18,22], whereas promoter knockout caused a substantial reduction in gene *10* transcript abundance [22]. Surprisingly, when combining the double promoter knockout Δ*ϕ*9/10 with codon deoptimization, fitness was nearly identical to the case of just the promoter knockout [22]. Thus, there were strong diminishing returns: The combined fitness-reducing effects of the two modifications were weaker than those of the individual modifications.

We attempted to recapitulate these result with our simulation, by simulating four different strains of bacteriophage T7: wildtype, a strain with gene *10* codon-deoptimized (i.e. recoded), a strain with *ϕ*10 and *ϕ*9 knocked out, and a strain with both modifications. We measured simulated gene *10* protein abundance, which we then used to approximate fitness (see Methods). We found that our simulation broadly recapitulated the experimental findings of the individual attenuation strategies, even though it missed specific details (Fig. 9). In particular, we found that codon deoptimization had a much smaller effect on fitness than promoter knockout. However, when combining attenuation strategies, the recoding and promoter knockout were nearly additive in simulations but showed diminishing returns in experiments. This finding suggested that our model is missing the mechanism responsible for the antagonistic effects of combining attenuation strategies. In summary, our mechanistic simulations recapitulated the effects of single knockouts, but overestimated the effect of combining attenuation strategies.

**Figure 9:**
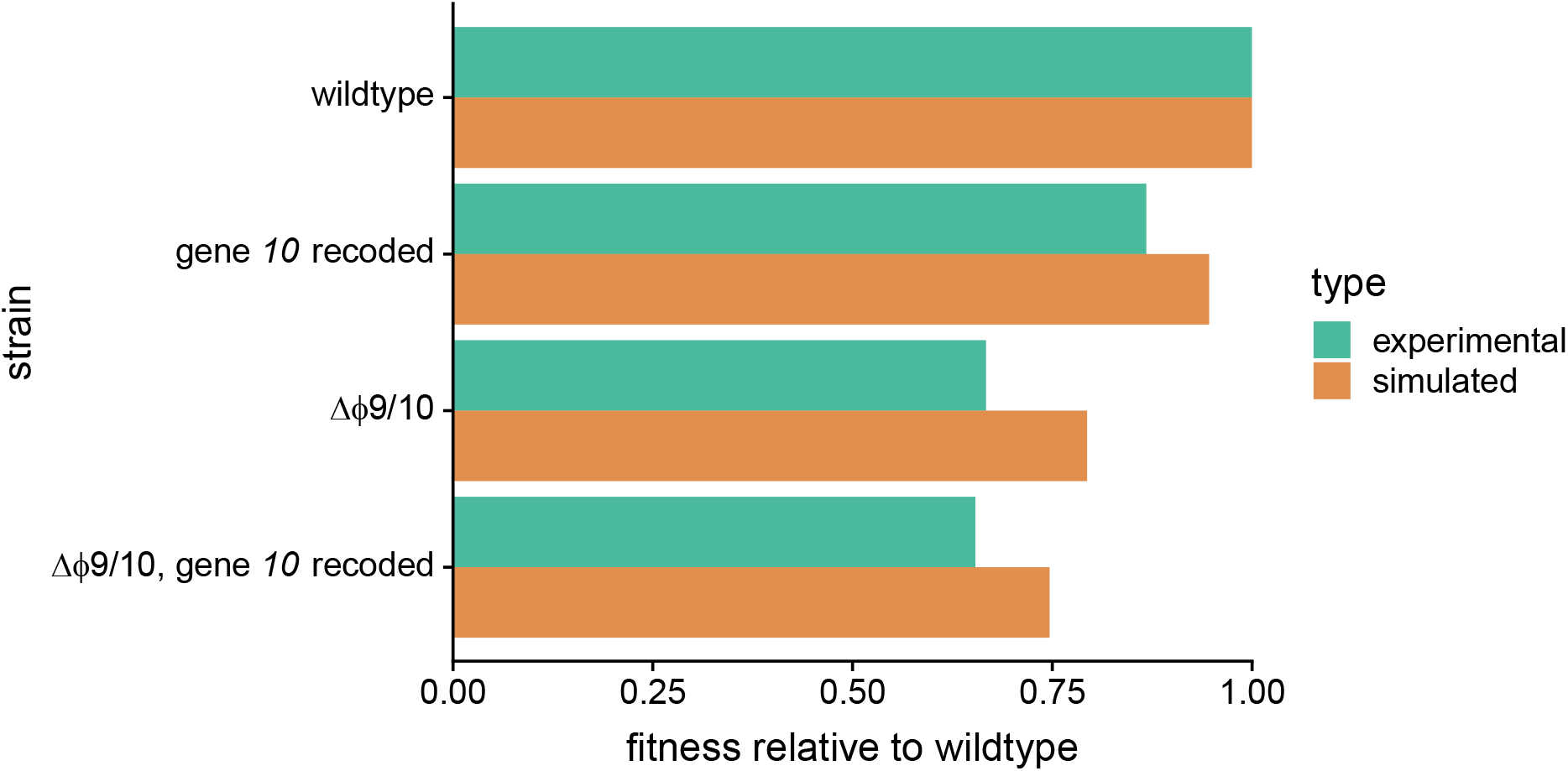
Predicted and experimental fitness of four different strains of T7, a wildtype strain, a strain in which gene *10* has been recoded to use non-preferred codons, a strain with the promoters *ϕ*9 and *ϕ*10 knocked out, and a strain with both the promoter knockouts and codon deoptimization. Fitness is shown relative to the wildtype strain. Simulations were conducted with transcript degradation. In the simulations, the recoding and promoter knockout reduced fitness by 5.4% and 20.6%, respectively, relative to wildtype. Combining these two modifications created a 25.3% reduction in fitness, nearly identical to the reduction expected from combining the individual effects (24.9%). By contrast, in the experiments the recoding and promoter knockout reduced fitness by 13.2% and 33.3%, respectively, but the genome with both modifications had a fitness reduction nearly identical to the promoter knockout alone, 34.6%. This difference suggests that our model does not fully capture the antagonistic effects of mixing attenuation strategies.

## Discussion

Advances in high-throughput RNA sequencing and mass spectrometry-based proteomics have recently allowed for genome-wide measurements of transcript and protein abundances over time in a growing bacteriophage T7 [18,22]. Here, we reanalyzed data collected from these two prior studies, in combination with newly measured RNA abundances for a non-lab-adapted strain T7^+^. The T7 genome is split into three classes [10]. We observed evidence for differential gene regulation among these classes, in both lab-adapted and non-lab-adapted strains. We also detected a set of down-regulated genes *(3.8–6.5)* within class II. We hypothesized that targeted transcript degradation caused this down-regulation, and validated this hypothesis using stochastic simulations of bacteriophage T7 gene expression. Our simulations of T7 that included directional transcript degradation and RNase cleavage sites recapitulated this down-regulated set of genes, and more broadly gene expression trends not explained by promoters and terminators, whereas our simulations of T7 without transcript degradation failed to capture these regulatory patterns. We next assessed the relationship between proteins and transcripts in both the experimental data and *in silico*. Here, we observed evidence for both differential gene expression among the three classes, and differences in correlations between protein and transcripts. Finally, we recreated several prior viral attenuation experiments [22] in our simulation. This simulation captured some aspects of the experiments but missed others. In particular, it could not reproduce the observation that combining different attenuation strategies produces diminishing fitness effects. In summary, we demonstrate evidence for extensive degradation of the T7 transcriptome. Moreover, simulations can provide mechanistic insight into these and other experimental findings, but need further refinements to make highly accurate predictions.

Bacteria use transcript degradation extensively as a strategy for gene regulation [27–32,37–39]. Degradation helps to tightly couple transcription and translation, such that protein abundances closely match those of transcripts and bacteria can more quickly respond to changes in their environment. In many cases, however, specific transcript properties drive differential degradation patterns [30]. In *E. coli* operons, for example, secondary structure creates patterns of differential degradation within single polycistronic transcripts [39]. Bacteriophage T7, which infects *E. coli,* produces almost exclusively polycistronic transcripts, which are processed by *E. coli* RNase III at specific cleavage sites [10,40,41]. However, T7 can grow in strains of *E. coli* lacking RNase III [37], and given the high stability of T7 transcripts [40], the broader role, if any, of transcript degradation in T7 is unclear. We emphasize that our 5’-to-3’ model of degradation is an approximation of the effects of multiple endo- and exonucleases in *E. coli*. In the most common pathway for transcript degradation, *E. coli* endonucleases first cleave the transcript in to small fragments, and then an exonuclease degrades the small fragments in the 3’-to-5’ direction [28]. This first step of cleavage tends to happen closer to the 5’-end of the transcript than to the 3’-end, and so the 5’-ends of transcripts generally degrade sooner [31]. Future versions of the Pinetree simulator could explicitly model both of these steps instead of approximating them with a 5’-to-3’ RNase. Regardless of these minor modeling choices, our findings suggest that extensive transcript degradation alters the distribution of transcripts in T7. Since T7 RNA polymerase synthesizes transcripts at a rate approximately 8 times faster than ribosomes translate them, transcription and translation are largely assumed to be uncoupled in T7 [34]. Controlled degradation of T7 transcripts may allow T7 to selectively recouple transcription to translation.

Our simulations of T7 gene expression made reasonable predictions for the effects of specific genomic modifications. In particular, our simulations predicted that promoter knockout had a much bigger effect on fitness than codon deoptimization, even if the exact fitness reductions were not accurately predicted. We note, however, that more quantitatively accurate predictions could be achieved via careful optimization of the various simulation parameters, which we did not undertake in this work.

Our simulations did not, however, correctly predict the reduced fitness effect of combining promoter knockout with codon deoptimization. Similarly, our simulations could not explain the weaker correlation that we observe between protein and transcript abundances in the experimental data. We believe that both discrepancies may be due to one important shortcoming of Pinetree, namely that it does not explicitly model tRNA pools. Our current simulations assume that the cell has unlimited protein production resources. However, we know that the availability of tRNAs plays an important role in translation and will influence the relative relationship between proteins and transcripts under different growth conditions [42–44]. Given the short life cycle of a T7 infection, the translation of phage proteins likely becomes limited by the availability of charged tRNAs. Future versions of Pinetree could explicitly incorporate these tRNA pools, similar to other stochastic models [43,45], and parameterize them by either ribosome footprinting or direct tRNA measurements [46,47]. In either case, the limitations of the current simulation point towards specific biological mechanisms that may affect T7 RNA and protein expression under certain conditions. Thus, our present work suggests both future improvements in the simulation and future experimental work.

Our T7 simulator is the next step forward in a long list of computational models of bacteriophage T7. The first models were simple kinetic models based on differential equations [12,13], and more recently coupled with flux-balance equations to describe the metabolism of the host [15]. For a recent review of this type of modeling, see Ref. [16]. Kinetic models can capture complex gene regulatory behavior and exhibit rich dynamics, but ultimately they are too simplistic to accurately describe gene expression. For example, the time to first production of a protein tends to be too small in kinetic models, because they don’t accurately capture the time it takes for a polymerase to process an entire transcript [14]. More realistic models track the movement of individual polymerases or ribosomes along DNA or RNA, using a stochastic framework. The first genome-scale model of this type was the stochastic gene expression simulator TABASCO [14], developed to describe T7 gene expression. TABASCO contains a stepwise transcription model, tracking the movement of individual polymerases along the phage genome. However, it treats translation via a kinetic model, and it does not contain a stepwise, directional degradation model. With our simulator Pinetree [26], we have built on the logic developed for TABASCO and have extended it to include stochastic, stepwise descriptions of both translation and transcript degradation. Our results here show that this is a viable pathway towards a realistic and efficient computational model of T7 gene expression.

As both computational modeling and experimental techniques become more sophisticated, we are approaching a point where models can inform experiments, test mechanistic hypotheses *in silico,* and make predictions of gene expression in highly dynamic environments. These advanced models will allow us to predict mutants with desired phenotypes, design viral genomes that are attenuated and/or display limited potential for adaptation, and generally unlock new engineering possibilities in synthetic biology.

## Methods

### RNA-sequencing analysis

We analyzed RNA-sequencing data from *E. coli* infected with T7 grown at 37 °C, and collected at 1, 5, and 9 minutes after infection [18]. We first created a reference sequence containing both the T7 (NCBI: NC_001604.1) and *E. coli* K12 (NCBI: U00096.3) genomes. As described previously [18], we excluded *10B* from our analyses, because it is a readthrough product of gene *10A* and most reads that map to *10B* will also map to *10A.* To simplify our notation, we refer to genes *10A* and *10B* jointly as gene *10.* We used HISAT2 to map the reads to our reference genome [48]. We generated raw read counts with BEDtools using the “multicov” command [49]. Lastly, we removed *E. coli* reads and converted raw read counts to transcripts per million (TPM) to get transcript abundance estimates [50]. To visualize individually mapped reads, we used the BEDtools “genomecov” command [49].

To obtain RNA abundances for the 30 °C time course, we isolated RNA from T7^+^-infected BL21 *E. coli* samples; T7^+^ was added at a multiplicity of infection (MOI) between 2.5 and 5.0 to a 10 mL culture of cells growing exponentially at 30 °C. At 5, 10, 15, 20, and 25 minutes post-infection, two 2 mL samples of phage-infected culture were collected and pelleted in a microcentrifuge. RNA isolation, library preparation, and sequencing were carried out as previously described [18]. In brief, RNA was isolated using Trizol (Invitrogen) reagent, following the manufacturer’s protocol. Library preparation and sequencing was performed by the University of Texas Genome Sequencing and Analysis Facility (UT GSAF). RNA samples were analyzed on an Agilent 2100 BioAnalyzer and libraries were prepared using the NEBNext Ultra II Library Prep Kit series. Sequencing was conducted on an Illumina HiSeq 2500 (SR50). Subsequent analysis was performed as described in the preceding paragraph.

### Proteomics data

We acquired processed proteomics data from the same study as the RNA-sequencing data [18]. These data include estimates of protein abundances from the same time points (1, 5, and 9 minutes post-infection), collected under the same conditions as the RNA-sequencing data. We made no modifications to the previously used analysis pipeline [18].

### Simulation models of three-gene plasmids

We constructed two linear plasmid models from which to simulate gene expression using Pinetree [26]. Each plasmid contained three genes (*X, Y*, and *Z*), each 150bp in length. We defined a single promoter upstream of gene *X* and a single terminator downstream of gene *Z*. For the second plasmid model, we added a second promoter immediately upstream of gene *Z* followed by an RNase cleavage site. We simulated gene expression for 240s in an *E. coli*-like environment at a reduced scale. The cell volume was 8 × 10^-16^ L, with initial conditions of 10 RNA polymerases and 100 ribosomes. Promoter strengths and rates of transcript cleavage and degradation were defined arbitrarily. Full parameter files for each simulation are available on GitHub and are archived on Zenodo (see *Code and data availability*).

### Simulation models of bacteriophage T7

To simulate bacteriophage T7 infecting *E. coli,* we again used Pinetree [26]. We constructed models with and without degradation. All models have the same initial conditions and parameters, except where noted below. Full parameter files for each simulation are available on GitHub and are archived on Zenodo (see *Code and data availability*).

#### Initial conditions and species-level reactions

The Pinetree simulator models transcription and translation at single-base resolution, but otherwise only supports pooled species-level reactions. These reactions model molecular species with specific copy numbers. The simulation assumes that the molecular species interact stochastically as described by the Gillespie algorithm [51]. For our model of T7, most of these species-level reactions were derived from a prior stochastic model of T7 (Tables 1 and 2) [14]. To more accurately account for the *E. coli* cellular resources available to T7 upon infection, we added several reactions. These reactions include the degradation of the *E. coli* genome, production of *E. coli* transcripts, and the binding and unbinding of ribosomes to *E. coli* transcripts. These *E. coli* transcripts differ from the T7 transcripts in that they are modeled at the species-level and not at the single-base level. All rate constants in these additional reactions were defined arbitrarily to conform to experimental transcript and protein distributions. Our aim was to approximate *E. coli* genome and transcript degradation and the shift in ribosomal resources towards the production of T7 proteins.

**Table 1:**
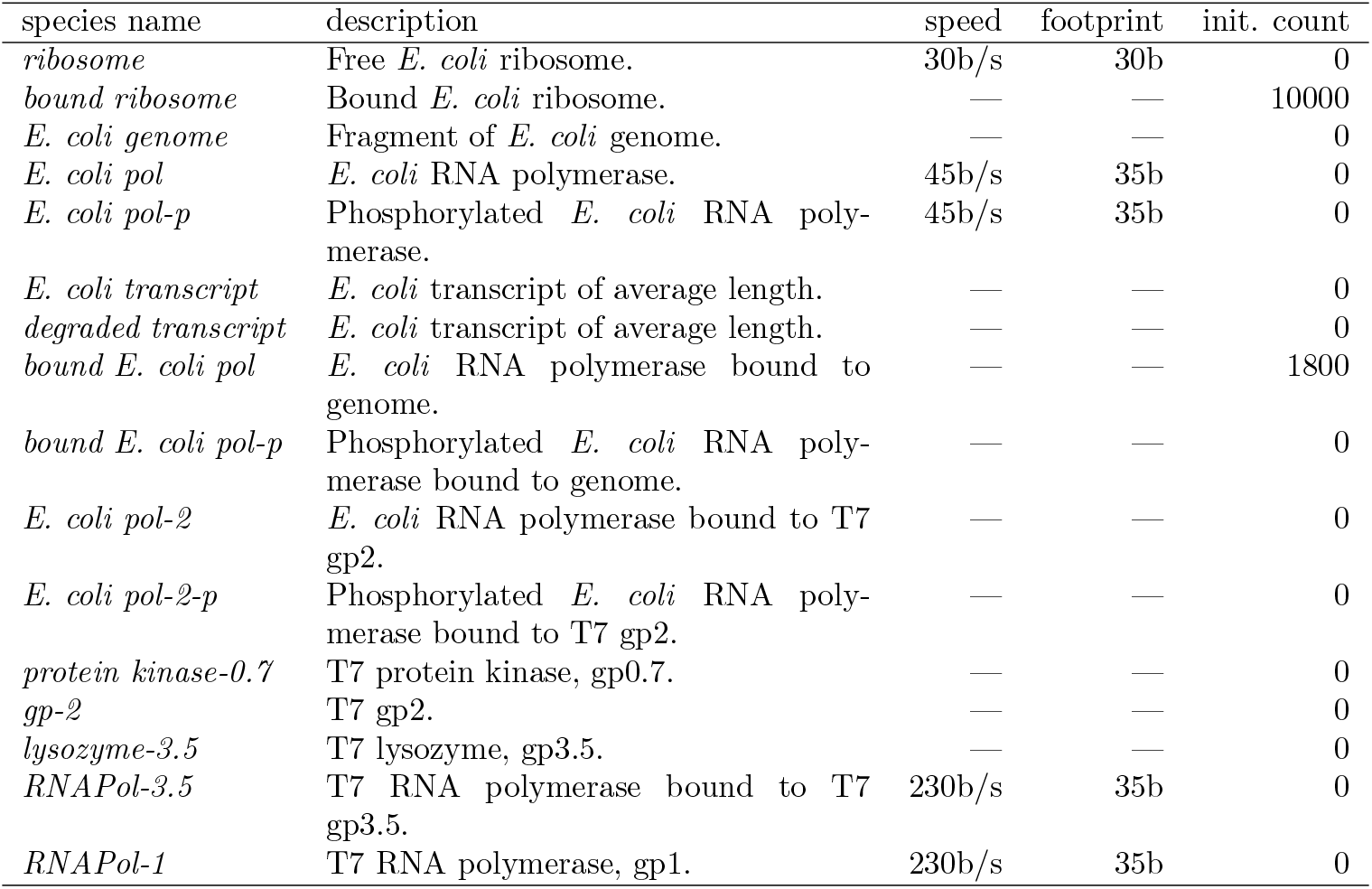
Molecular species in simulations of bacteriophage T7 gene expression.

| species name | description | speed | footprint | init. count |
| --- | --- | --- | --- | --- |
| *ribosome* | Free *E. coli* ribosome. | 30b/s | 30b | 0 |
| *bound ribosome* | Bound *E. coli* ribosome. | — | — | 10000 |
| *E. coli genome* | Fragment of *E. coli* genome. | — | — | 0 |
| *E. coli pol* | *E. coli* RNA polymerase. | 45b/s | 35b | 0 |
| *E. coli pol-p* | Phosphorylated *E. coli* RNA polymerase. | 45b/s | 35b | 0 |
| *E. coli transcript* | *E. coli* transcript of average length. | — | — | 0 |
| *degraded transcript* | *E. coli* transcript of average length. | — | — | 0 |
| *bound E. coli pol* | *E. coli* RNA polymerase bound to genome. | — | — | 1800 |
| *bound E. coli pol-p* | Phosphorylated *E. coli* RNA polymerase bound to genome. | — | — | 0 |
| *E. coli pol-2* | *E. coli* RNA polymerase bound to T7 gp2. | — | — | 0 |
| *E. coli pol-2-p* | Phosphorylated *E. coli* RNA polymerase bound to T7 gp2. | — | — | 0 |
| *protein kinase-0.7* | T7 protein kinase, gp0.7. | — | — | 0 |
| *gp-2* | T7 gp2. | — | — | 0 |
| *lysozyme-3.5* | T7 lysozyme, gp3.5. | — | — | 0 |
| *RNAPol-3.5* | T7 RNA polymerase bound to T7 gp3.5. | 230b/s | 35b | 0 |
| *RNAPol-1* | T7 RNA polymerase, gp1. | 230b/s | 35b | 0 |

**Table 2:** Species-level reactions and rate constants used in simulations of bacteriophage T7 gene expression.

| reaction | rate constant |
| --- | --- |
| *ribosome* + *E. coli transcript* → *bound ribosome* | $10^6 \text{ M}^{-1}\text{s}^{-1}$ |
| *bound ribosome* → *ribosome* + *E. coli transcript* | $4 \times 10^{-2} \text{ s}^{-1}$ |
| *E. coli transcript* → *degraded transcript* | $1.925 \times 10^{-3} \text{ s}^{-1}$ |
| *E. coli pol* + *E. coli genome* → *bound E. coli pol* | $10^7 \text{ M}^{-1}\text{s}^{-1}$ |
| *E. coli pol-p* + *E. coli genome* → *bound E. coli pol-p* | $3 \times 10^6 \text{ M}^{-1}\text{s}^{-1}$ |
| *bound E. coli pol* → *E. coli transcript* + *E. coli genome* + *E. coli pol* | $4 \times 10^{-2} \text{ s}^{-1}$ |
| *bound E. coli pol-p* → *E. coli transcript* + *E. coli genome* + *E. coli pol-p* | $4 \times 10^{-2} \text{ s}^{-1}$ |
| * *protein kinase-0.7* + *E. coli pol* → *protein kinase-0.7* + *E. coli pol-p* | $3.8 \times 10^7 \text{ M}^{-1}\text{s}^{-1}$ |
| * *protein kinase-0.7* + *E. coli pol-2* → *protein kinase-0.7* + *E. coli pol-2-p* | $3.8 \times 10^7 \text{ M}^{-1}\text{s}^{-1}$ |
| * *gp-2* + *E. coli pol* → *E. coli pol-2* | $3.8 \times 10^7 \text{ M}^{-1}\text{s}^{-1}$ |
| * *gp-2* + *E. coli pol-p* → *E. coli pol-2-p* | $3.8 \times 10^7 \text{ M}^{-1}\text{s}^{-1}$ |
| * *E. coli pol-2* → *gp-2* + *E. coli pol* | $1.1 \text{ s}^{-1}$ |
| * *E. coli pol-2-p* → *gp-2* + *E. coli pol-p* | $1.1 \text{ s}^{-1}$ |
| * *lysozyme-3.5* + *RNAPol-1* → *RNAPol-3.5* | $3.8 \times 10^9 \text{ M}^{-1}\text{s}^{-1}$ |
| * *RNAPol-3.5* → *lysozyme-3.5* + *RNAPol-1* | $3.5 \text{ s}^{-1}$ |

\* Derived from Kosuri, et al [14].

Each simulation begins with 500bp of the T7 genome already injected into an *E. coli* cell. The cell has a volume of 1.1 × 10^-15^ L. Initially, free *E. coli* RNA polymerases bind to the early T7 promoters. Once the T7 RNA Polymerase has been translated, it begins transcribing later T7 genes at 230bp/s and pulls the remainder of the genome into the cell [20]. We assume that a single phage genome infects a single *E. coli* cell. Upon infection, we also assume that the *E. coli* cell contains 10,000 actively translating ribosomes *(bound ribosome*) and 1800 *E. coli* RNA polymerases *(E. coli RNA pol*). These quantities were derived from Kosuri, et al [14]. We do not explicitly model T7 genome replication, and we assume that all gene expression occurs from a single T7 genome.

#### Promoter, terminators, and genome organization

All genes and regulatory elements in our models of T7 were generated directly from the annotated genome (NCBI: NC_001604.1). We included all genes except for genes *0.4, 0.5, 0.6A, 0.6B, 5.5-5.7, 4.1, 4B, 10B.* These genes were excluded because either they were not present in the strains of T7 used in our experiments or because of limitations in Pinetree. For example, Pinetree does not support translational readthrough products such as the minor capsid protein encoded by gene *10B.*

We included all promoters in T7, except for weak promoters near the origin of replication *(E. coli* promoters A0 and E6, and T7 promoters *ϕ*OR and *ϕ*OL were excluded). Promoter strengths are defined relative to the strongest promoter, *ϕ*10 (Table 3). We derived these relative promoter strengths from a prior deterministic model of bacteriophage T7 infection [15]. Promoters themselves are defined as 35bp regions of the genome directly upstream of the transcription start site in the reference genome. This 35bp length is the footprint of all RNA polymerases. In Pinetree, all promoters must be at least as long as the footprint of the polymerases that bind to them. Some promoter strengths were modified to better fit the distribution of transcript abundances observed in our experimental data. For simulations without degradation, we set the *ϕ*10 promoter strength to 1.82 × 10^7^ M^-1^s^-1^ [15]. We arbitrarily increased this rate constant to 1.82 × 10^8^ M^-1^s^-1^ for simulations with degradation to maintain similar absolute transcript abundances between simulations with and without degradation. In simulations of promoter knock-out strains, we set the promoter strengths of the knocked-out promoters to zero.

**Table 3:** Promoter strengths in simulations of T7 gene expression. These are promoter strengths for *E. coli* polymerases that are unposphorylated and unbound to gp2, and for T7 RNA polymerase unbound to lysozyme.

| promoter | strength |
| --- | --- |
| *E. coli* promoter A1 | $10^5 \text{ M}^{-1}\text{s}^{-1}$ |
| *E. coli* promoter A2 | $10^5 \text{ M}^{-1}\text{s}^{-1}$ |
| *E. coli* promoter A3 | $10^5 \text{ M}^{-1}\text{s}^{-1}$ |
| *E. coli* promoter B | $10^4 \text{ M}^{-1}\text{s}^{-1}$ |
| *E. coli* promoter C | $10^4 \text{ M}^{-1}\text{s}^{-1}$ |
| ϕ1.1A | 0.01* |
| ϕ1.1B | 0.01* |
| ϕ1.3 | 0.01* |
| ϕ1.5 | 0.01* |
| ϕ1.6 | 0.01* |
| ϕ2.5 | 0.01* |
| ϕ3.8 | 0.01* |
| ϕ4C | 0.01* |
| ϕ4.3 | 0.01* |
| ϕ4.7 | 0.01* |
| ϕ6.5 | 0.05* |
| ϕ9 | 0.01* |
| ϕ10 | $1.82 \times 10^7 (1.82 \times 10^8)^\dagger \text{ M}^{-1}\text{s}^{-1}$ |
| ϕ13 | 0.1* |

\* Promoter strength relative to *phi*10.
<sup>†</sup> Promoter strength for simulations with degradation, which was increased so that absolute transcript abundances were comparable among simulations with and without degradation.

In bacteriophage T7, gene *3.5* lysozyme facilitates the transition from expression of class II genes to class III genes [10,11]. To simulate this transition, we modeled gene *1* T7 RNA polymerase bound to lysozyme and unbound to lysozyme as separate molecular species. Employing a rate constant from prior stochastic T7 simulations [14], lysozyme and T7 RNA polymerase bind to form this new gene *1–3.5* complex (Table 2). These two polymerases have different binding affinities to promoters. For all class II promoters, the *1–3.5* complex binds with a rate constant of 0.5 times that of the rate with only T7 RNA polymerase. During the simulation, as abundances of lysozyme increase, this differential binding interaction has the effect of shifting promoter binding preferences from class II to class III.

For the regulation of *E. coli* RNA polymerase, we defined a set of reactions similar to that of T7 RNA polymerase regulation (Table 2). Gene *0.7* kinase posphorylates *E. coli* RNA polymerase, modulating its binding activity. Gene *2* reacts with *E. coli* RNA polymerase and deactivates it entirely. Rate constants for these reactions were derived from Kosuri, et al [14]. Together, these reactions impede the interaction between *E. coli* RNA polymerase and the *E. coli* promoters within the early region of the T7 genome. As the simulation progresses, transcription from these early promoters becomes negligible.

#### Ribosome binding sites and translation

Ribosome binding site strengths were derived from a prior stochastic simulation of T7 [14]. We defined ribosome binding sites as the 30 bp regions immediately upstream of start codons. Again, this definition is due to a limitation in Pinetree, where the binding site region must be at least as large as the footprint of the ribosome. Ribosomes move step-wise along the mRNA at an average rate of 30 bp/s, which can be scaled up or down depending on the position within the transcript. We used this scaling factor to simulate codon deoptimization. In the simulations of T7 with codon-deoptimized gene *10,* we scaled the translation rate of all codons within gene *10* by a factor of 0.2.

#### Degradation model

We employed a directional model of transcript degradation parameterized by two different initiation rate constants and a degradation rate. We assumed that transcripts degrade in the 5’-to-3’ direction. We note that there are no 5’-to-3’ exonucleases in *E. coli* but that degradation occurs via several different endo- and exonucleases [28]. On average, however, the 5’-end of transcripts tend to have a shorter half-life than the 3’-end [31]. To simulate this directional degradation effect in a way that was computationally tractable, we implemented a 5’-to-3’ exonuclease in Pinetree. Internally, Pinetree appends an RNase binding site of 10 bp in length to the 5’ end of each newly-synthesized transcript. To represent RNase cleavage sites, we defined an RNase binding site at each cleavage site. These two types of binding sites differ in their binding rate constants: We assumed a rate constant of 10^-2^ s^-1^ for cleavage sites and 10^-5^ s^-1^ for 5’-end sites. Although these absolute rate constants were defined arbitrarily, we assigned a lower rate 5’-end binding sites because of additional phosphate cleavage steps that occur before degradation begins [27,32]. The binding reaction itself is unimolecular and depends only on the abundance of the transcripts. Once an RNase has bound, it degrades transcripts in the 5’-3’ direction at a rate of 20 bp/s, again defined arbitrarily to approximate the shorter lifespan of the 5’-end of transcripts [31].

### Comparing simulations to experiments

The rate constants in our simulations were originally derived from experiments conducted at either 25 °C or 30 °C [14,15]. At these temperatures, T7 has a lysis time of approximately 20-25 minutes [10,11]. In contrast, much of the experimental data we analyzed had been collected at 37 °C [18], when the phage lyse at 11 minutes. Thus, 9 minutes in our simulations is not directly comparable to 9 minutes in the experimental data. To compare simulated and experimental gene expression, we selected 500s and 1000s in the simulation to represent 5 minutes and 9 minutes, respectively, in the experimental data taken at at 37 °C.

To calculate simulated fitness in doublings per hour, we made use of a previously published relationship between intrinsic growth rate (*r*) and burst size (b) [17]:

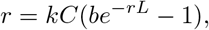

where *k* is adsorption rate, *C* is cell density, and *L* is lysis time. We assumed lysis time, adsorption rate, and cell density are all fixed constants. We set *L* to 12 minutes and arbitrarily set *kC* to 1. To estimate burst size, we assumed that all capsid proteins present at 1000s are converted into virions, and that there are 400 copies of the capsid protein per virion. Using these assumptions, we arrived at the following equation relating the intrinsic growth rate *r* to simulated capsid protein counts *p*:

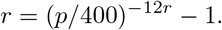

We solved numerically for *r* numerically using Wolfram Alpha for each of the four simulated conditions: wildtype, gene 10A recoded, phi9/10 double knockout, and the double knockout combined with the recoding. We converted intrinsic growth rate *r* to doublings per hour *d* using the following equation:

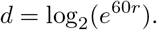

Lastly, we normalized all fitness values to that of the wildtype.

### Code and data availability

All code and processed data used to produce the figures and analyses presented here is available on Github (https://github.com/benjaminjack/phage_simulation) and is archived on Zenodo (DOI: 10.5281/zenodo.2631365). This archive includes specific parameter files for all simulations described in this work. Raw sequencing reads for the 30 °C time course have been submitted to the NCBI Gene Expression Omnibus under accession number GSE123854.

## Acknowledgments

This work was supported by National Institutes of Health grant R01 GM088344. We thank Jim Bull for feedback and use of lab space for the collection of samples. We thank Ian Molineux for providing us with an isolate of T7^+^ and for suggesting that transcript degradation is a critical component of gene regulation in T7.

